# Pharmacodynamic and stage-dependent therapeutic efficacy of SFRP1 neutralization in a mouse model of Alzheimer’s disease

**DOI:** 10.64898/2026.03.26.714543

**Authors:** Pablo Miaja, Marcos Martinez-Baños, María Jesús Martin-Bermejo, Inmaculada Moreno, Mercedes Dominguez, Paola Bovolenta

## Abstract

Alzheimer’s disease (AD) is characterized by early synaptic dysfunction followed by progressive amyloid-β (Aβ) accumulation, neuroinflammation, and cognitive decline. We previously identified Secreted Frizzled-Related Protein 1 (SFRP1) as a multifactorial contributor to AD pathogenesis and provided initial evidence that its neutralization ameliorates pathological AD-like traits in mice. Here, we evaluate the pharmacodynamics, biodistribution, and therapeutic window of an α-SFRP1 monoclonal antibody (α-SFRP1) in APP/PS1 mice. Pharmacokinetics and target engagement of α-SFRP1 were assessed in different groups of APP/PS1 mice using biotinylated or ^89^Zr-labelled antibodies, with tissue distribution and α-SFRP1 levels quantified by in-house ELISA or PET/CT. Therapeutic efficacy was evaluated by administering α-SFRP1 or the SFRP1 inhibitor WAY-316606 at different stages of disease progression via retro-orbital injection, followed by analysis of AD-like pathology using ELISA and immunofluorescence assays followed by quantifications and statistical analysis. Using ^89^Zr-labelled antibodies, we show that intravenously administered α-SFRP1 engages its target systemically and reaches the brain, although at substantially lower levels and with a rapid 24-hour clearance. Treatment with α-SFRP1 had no apparent systemic side effects, but its therapeutic efficacy against AD-like brain pathology was strongly dependent on disease stage. While early administration reduced amyloid pathology in previous studies, treatment initiated at intermediate or advanced stages showed minimal benefit at standard doses. Higher antibody doses reduced amyloid burden and dystrophic neurites but were associated with increased mortality with time. Pharmacological inhibition of SFRP1 using a small-molecule inhibitor similarly failed to ameliorate pathology at intermediate stages. Together, these findings demonstrate that SFRP1 remains a relevant therapeutic target in AD, but its effective modulation is constrained by limited brain exposure and a narrow therapeutic window, underscoring the importance of early intervention and prompting the search for improved brain-targeted delivery strategies.

## 1. Introduction

Neurodegenerative diseases (NDs) comprise a variety of progressive and devastating brain disorders that primarily affect the elderly, thereby representing a major socio-economical and emotional burden for our increasingly aging population. Alzheimer’s Disease (AD) represents around 80% of the ND cases, making it the most prevalent form of dementia worldwide (1). AD was first clinically described as an “unusual illness of the cerebral cortex” in 1907 (2). Since then, enormous research efforts have been undertaken to understand its origin and pathophysiological mechanisms.

AD is considered a multifactorial disease characterized by synaptic loss, the accumulation of extracellular and vascular amyloid-β (Aβ) containing plaques, the formation of intracellular hyperphosphorylated tau tangles, a chronic state of glia-driven neuroinflammation, vascular and neuronal damage and an overall brain atrophy, finally leading to cognitive impairment (3). Despite the large amount of published studies and the numerous hypotheses on the disease causes (4–6) the trigger(s) for disease onset remain unclear. Early synaptic loss and chronic neuroinflammation are considered key events of the disease (7,8). Nevertheless, the deposition of toxic Aβ aggregates in the brain is recognized as a main hallmark of AD, representing the basis of the so-called amyloid-cascade hypothesis (9,10).

Aβ peptides originate from the proteolytic processing of the Amyloid Precursor Protein (APP), which is heterogeneously expressed throughout the body with particularly high levels in brain neurons (11). APP can be processed via two physiological pathways, resulting in the release of different soluble fragments: the amyloidogenic (producing Aβ monomers) and the non-amyloidogenic pathways. In both cases, APP undergoes a two-steps proteolytic cleavage. In the non-amyloidogenic pathway, A Disintegrin and Metalloproteinase domain-containing protein 10 (ADAM10) cleaves APP within the Aβ domain preventing the formation of Aβ peptides upon presenilins 1 and 2 (PSEN1/2) cleavage (12). When escaping this processing, APP undergoes amyloidogenic cleavage via the β-site APP cleaving enzyme 1 (BACE1) followed by PSEN1/2. If the balance between these pathways favours the generation and accumulation of Aβ peptides, amyloid plaques build up in the brain, constituting a diagnostic AD hallmark.

The widespread recognition of this hallmark has led to the development of therapeutic monoclonal antibodies (mAb) against Aβ as the first disease-modifying therapies for AD (13). Nevertheless, this approach is under debate (14) and only two of these mAb immunotherapies (Dodanemab and Lecanemab) are currently being commercialized in a limited number of countries (15,16), highlighting the need of exploring new therapeutic targets.

Previous work from our lab identified Secreted Frizzled Related Protein 1 (SFRP1) as a novel multifactorial player in AD pathogenesis (17). SFRP1 is a small (35 kDa) soluble protein that modulates Wnt signalling (18) and regulates the enzymatic activity of ADAM10 (19). Our pioneer studies showed that SFRP1 abnormally accumulates in the brain of AD patients and APP/PS1 mice (17). This finding has thereafter been supported by a series of proteomic studies on large patient cohorts, consistently confirming upregulated SFRP1 levels in the frontal cortex and hippocampus from AD patients (20–27) and related mouse models (23,28). Functional studies have shown that SFRP1 not only favours Aβ formation and accumulation by interfering with ADAM10-mediated APP processing, but also promotes the presence of toxic Aβ peptides, deposits in amyloid plaques (17) and sustains a chronic neuroinflammatory response (29). Additionally, chronic exposure to SFRP1 alone is sufficient to induce synaptic deficits and cognitive impairments resembling the neurodegenerative progression observed in AD (30). Most notably, genetic or mAb-mediated neutralization of SFRP1 reduce amyloid plaque burden and prevents the loss of LTP response commonly observed in APP/PS1 mice (17).

Together with the current need for a broader molecular and mechanistic understanding of AD (31), these findings support the therapeutic potential of targeting SFRP1 in AD. Here, we have further explored the therapeutic efficacy of the already described α-SFRP1.10.5.6 mAb (α-SFRP1, in brief) (17) together with its pharmacodynamics. We report that intravenous (IV) administration of the mAb enables target engagement in the brain, although at much lower levels than in other organs. mAb administration has no apparent side effects but its therapeutic window is limited to early stages of the disease.

## 2. Methods

### 2.1. Mice

Throughout the study, double chimeric transgenic APP/PS1 mice served as experimental model. These mice express the human mutated APP (APP^695swe^) and presinilin1 (PS1^dE9^) genes under the prion promoter, which directs expression to the central nervous system (CNS) (32). Male and female homozygous (APP/PS1^Hom^) and heterozygous (APP/PS1^Het^) mice aged 4–12 months were included, based solely on genotype and age as inclusion/exclusion criteria, with sex-balanced group allocation whenever possible. Animals were housed at the animal facility of the Centro de Biología Molecular Severo Ochoa (CBM) or in the Centro de Investigación Cooperativa en Biomateriales (CIC biomaGUNE) for radiolabeled antibodies administration. In both cases, animals were housed according to European Communities Council Directive of 24 November 1986 (86/609/EEC); in a temperature-controlled, pathogen-free environment under 12 h light-dark cycles, and available food and water *ad libitum*. All procedures were approved by the ethical committees of the CBM, the Bioethics sub-committee of Consejo Superior de Investigaciones Científica (CSIC, Madrid, Spain) and the Comunidad Autónoma de Madrid under the protocol approval number PROEX 092.6/21; RD 53/2013 or the Ethical Committee of Centro de Investigación Cooperativa en Biomateriales (CIC biomaGUNE) and authorized by the local Diputación Foral de Gipuzkua under the protocol approval number PRO-AE-SS-212 in case of radiolabeled antibodies administration.

#### 2.2. Generation and biotinylation of α-SFRP1 monoclonal antibodies

Procedures for mAb generation were approved by the Instituto de Salud Carlos III (ISCIII) ethical committee (CBA PA 73-2011). α-SFRP1 antibodies were generated and characterized as detailed in (17). α-SFRP1.10.5.6 (α-SFRP1) and IgG1 antibodies were biotinylated using the EZ-link^TM^ NHS-PEG4-Biotinylation Kit (21455, Thermo Fisher, USA) according to manufacturer’s protocol. Briefly, each antibody was incubated at a 20-fold molar excess of biotin reagent (5 mg/mL in PBS) during 1h at RT. Biotin excess was removed by using a desalting column equilibrated with PBS and centrifuged at 1000 g for 2 min.

#### 2.3. Biodistribution of ^89^Zr-labeled antibodies by Positron Emission Tomography

[^89^Zr]Zr oxalate was purchased from Perkin Elmer whereas all other reagents used to label the mAbs were purchased from Sigma-Aldrich, USA, and used without further purification. Ultrapure water (resistivity > 18 MΩ cm) was generated using a Mili-Q system (Millipore, USA). The Radio-TLC from Raytest was used for the monitorization of the reaction as well as the stability tests. α-SFRP1 (2 mg/ml) and nonspecific IgG1 (2 mg/ml) bioconjugation with deferrioxamine-p-benzylisothiocyanate (DFO-Bz-NCS) and ^89^Zr-labeling was performed as previously described (33). Briefly, a 3-fold molar excess of DFO-Bz-NCS (5 mg/ml) was added to each antibody using 0.1M of sodium carbonate buffer (pH 9). The reaction mixture was incubated for 45 min at 37 °C, and the unreacted excess chelator was removed using the 40 kDa cutoff Zeba spin desalting columns (Thermo Fischer Scientific, MA, USA). Antibodies were then radiolabeled with ^89^Zr. Briefly, 50 µl of 1 M oxalic acid containing ^89^Zr ([^89^Zr]ZrC_2_O_4_; 0.6 mCi = 22.2 MBq) were neutralized with 1 M sodium carbonate. Then, 450 µg of either antibody was added and the volume adjusted to 0.5 ml with HEPES buffer (0.5 M). After 2.5 (α-SFRP1) or 6 (IgG1) hours of incubation at RT, the antibodies were purified by Sephadex G-25 size exclusion column (NAP5 GE-Healthcare) with PBS. Incubation and purification were monitored by iTLC (mobile phase: 20 mM citric acid xontaining 60 mM EDTA 9:1 acetonitrile; Rf (α-SFRP1/IgG1) = 0, Rf (^89^Zr-EDTA) = 0.8).

#### 2.4. Antibodies or drug administration

Mice divided in different groups and indicated in each specific case, were treated with either α-SFRP1 mAb or a non-specific IgG1 mAb as control (IgG1, MAB002, Bio-Techne, MN, USA), or with SFRP1 inhibitor WAY-316606 (WAY; S5815, Selleck Chemicals, USA) using, in this case, the DMSO diluent as control. These reagents were administered via intraperitoneal (IP) or retro-orbital sinus (RO) injections. Groups of animals were also injected via the tail vein with 100 µg (ca. 150 µl, 60 µCi) of radiolabeled antibody (α-SFRP1^89Zr^ or IgG1^89Zr^). RO and intra tail injections were performed in anesthetize mice (1.5-2% isoflurane in pure O_2;_ IsoFlo, Abbot Laboratories, USA or IsoVet, Spain). In the case of injections with radiolabeled antibody, animals were allowed to recover from anesthesia until defined scan time points, when animals were anesthetized again and monitored during the whole scan. In all cases, at the end of the treatment, animals were perfused and samples collected as described hereafter. In a first set of experiments, α-SFRP1 and IgG1 were labeled with biotin. A single dose of 100 µL of either of the antibodies at 1 µg/µL was injected via IP or RO in 4-8 months-old APP/PS1^hom^ mice for 24 h. The presence of biotinylated mAb was then determined by incubating 15 µm cryostat sections with streptavidin-POD at 1/500 followed by tyramide (Cell Signaling, USA). In a second set of experiments, several groups of homozygous mice of different ages were injected weekly via the RO with either α-SFRP1 or IgG1 as follows: 100 µl (1 mg/ml) were administered to 9 months-old mice for 3 months; to 4 months-old mice for 5 months; and 100 µl (2 mg/ml) were administered to 4 months-old mice for 2 months. The third group consisted of heterozygous mice that were treated with weekly RO injections of 100 µl of WAY (1 µM)-corresponding to slightly more than its in vitro calculated EC50 value (34) or 2% DMSO for 2 months starting at 8 months of age.

#### 2.5. PET/CT Imaging

Imaging studies were conducted using positron emission tomography (PET) in combination with computed tomography (CT), using the β and X-cube microsystem of Molecubes (MOLECUBES NV, Belgium). Static whole-body images (1 bed) were acquired in 511 keV ± 30% energetic window at 2, 10, 24, and 168 h post-administration (acquisition time = 45 min). CT acquisitions were also performed at the end of each PET scan, providing anatomical information for attenuation correction and for unambiguous localization of the radioactive signal. The reconstruction of the images was performed using the mathematic algorithm 3D Ordered Subset Expectation Maximization (3D-OSEM; 30 iterations).

#### 2.6. PET data analysis

PET images were analyzed using PMOD analysis software (PMOD Technologies Ltd., Switzerland). PET images were first co-registered with their corresponding CT reference. A standardized uptake value (SUV) correction factor (calculated based on the injected dose, the system’s internal calibration factor, and the animal’s body weight) was then applied to normalize voxel intensities to SUV units. Two distinct sets of volumes of interest (VOIs) were subsequently analyzed on the SUV-normalized PET images: (1) Body VOIs, which were manually delineated for individual organs (brain, lungs, heart, liver, kidneys, and bladder); and (2) Brain VOIs, which were defined using a mouse brain MRI atlas (M. Mirrione_T2, available within PMOD) to extract SUV values from specific brain regions. The CT image from each animal was co-registered to the MRI brain atlas using the CT–MRI transformation matrix (CT_MRI_coreg). This matrix was subsequently applied to align the corresponding PET image with the atlas, enabling the extraction of SUV values from specific brain regions. Results are presented as mean SUV ± SD from at least three animals.

#### 2.7. Samples collection

Blood samples were collected prior to the first injection and immediately before euthanasia. A 25 G needle was used to extract the blood from the submandibular vein. Blood was allowed to coagulate for 30 min at RT and then centrifuged for 30 min at 4°C and 1500g. The resulting serum was diluted 1:2 in PBS containing protease inhibitors and stored at −20 °C until use. At the end of the treatment, mice were anesthetized with ketamine/xilacine (Sigma), diluted each 1:10 in saline, and transcardiacally perfused with ice-cold 0.9% saline solution. Organs of interest were extracted and cut in equivalent halves. One half was post-fixed in 4% PFA for 24 h, washed in PBS and incubated in 30% sucrose solution in PBS for 48 h. Organs were embedded and frozen in a 7.5:15% gelatin:sucrose solution at -80°C until use. All organs were sectioned coronally at 15 µm thickness with a cryostat. The remaining half of each organ was used for protein extraction. Briefly, tissues were homogenized in RIPA buffer (700 µl) supplemented with protease inhibitors. Following mechanical homogenization, samples were incubated on ice for 30 min and then centrifuged at 21,000g for 30 min at 4 °C. The resulting supernatant, corresponding to the soluble protein fraction, was collected and stored at −80 °C until further use.

#### 2.8. Hematologic assessment

Blood samples, collected as explained above, and immediately mixed with EDTA (1 mg/mL of blood) to prevent clogging. Samples were kept in ice until analysis. Blood cell populations were quantified immediately using an Element HT5 veterinary hematology analyzer (Heska).

#### 2.9. ELISA assay for biotinylated antibody detection

Ninety-six-well microtiter plates (Nunc96, Thermo Fisher, USA) were coated overnight at 4°C with a goat α-mouse (1.5 µg/ml in PBS) using 50 µl/well. Plates were washed three times with PBST and incubated for 3h on a shaker with 100 µl of 2% BSA in PBST at RT, washed with PBST and incubated for 2 h at 37°C with 40 µl/well of serum diluted 1:2 in 2% BSA-PBST or 5 µg of brain lysates from mice injected with biotinylated α-SFRP1 or IgG1. Total protein concentration of the brain lysates was previously quantified by BCA Protein Assay Kit (Thermo Fisher, USA). Wells were washed with PBST and incubated for 1h at RT with streptavidin-POD (1:25000 in PBS, 50 µl/well; Jackson Laboratory, USA). After extensive washing with PBST, the enzymatic reaction was developed at RT in the dark for 20 min using tetra-methyl-benzidine liquid substrate (100 µl/well; Sigma-Aldrich, USA). The reaction was terminated by adding 2N HCl (100 µl/well), and the resulting product was measured at 450 nm in a microtiter plate ELISA reads FLUOstar OPTIMA (BMG Lab Tech, Germany).

#### 2.10. ELISA assay for SFRP1 detection

Ninety-six-well microtiter plates (Nunc96; Thermo Fisher, USA) were coated overnight at 4°C with α-SFRP1 (1.5 µg/ml in PBS; 50 µl/well). Plates were washed three times with PBST and incubated for 3 h on a shaker with 100 µl of 2% BSA in PBST at RT, washed with PBST and incubated for 1 h at 37°C with 40 µl/well of serum diluted 1:2 in 2% BSA-PBST or 5 µg of brain lysates. Total protein concentration of the brain lysates was previously quantified by BCA Protein Assay Kit (Thermo Fisher, USA). Wells were washed with PBST and incubated for 1h at 37°C with of biotin-labeled α-SFRP1.17.8.13 mAb (1 µg/ml in PBS; 50 µl/well) (17). Plates were further washed with PBST and incubated for 1 h at RT with streptavidin-POD (1:25000 in PBS; 50 µl/well Jackson Laboratory, USA). After extensive washing with PBST, the enzymatic reaction was developed at RT in the dark for 20 min using tetra-methyl-benzidine liquid substrate (100 µl/well; Sigma-Aldrich, USA). The reaction was terminated by adding 2N HCl (100 µl/well), and the resulting product was measured at 450 nm in a microtiter plate ELISA reads FLUOstar OPTIMA (BMG Lab Tech, Germany).

#### 2.11. Primary and secondary antibodies

The primary and secondary antibodies used in this study are listed in Table S1 indicating the source and the used dilutions.

**Table S1.** Immunofluorescence antibody information.

|  | Host | Dilution (IF) | Reference |
| --- | --- | --- | --- |
| **Fluorescent dye** |  |  |  |
| Thioflavine S | - | 0.01% | Sigma-Aldrich, 1326-12-1 |
| **Primary antibodies** |  |  |  |
| Aβ<sub>42</sub> (H31L21) | Rabbit | 1:500 | Invitrogen, 700254 |
| GFAP | Rabbit | 1:1000 | Dako, Z0334 |
| Iba1 | Rabbit | 1:500 | Wako, 019-19741 |
| LAMP1 (1D4B) | Mouse | 1:100 | DSHB, AB_528127 |
| **Secondary antibodies** |  |  |  |
| α-Mouse IgG 647 | Donkey | 1:1000 | Thermo Fischer, A-31571 |
| α-Rabbit IgG 488 | Donkey | 1:1000 | Thermo Fischer, A-31572 |
| α-Rabbit IgG 555 | Donkey | 1:1000 | Thermo Fischer, A-21206 |

#### 2.12. Immunofluorescence assays

Cryostat sections (15 µm) were allowed to thaw and dry and then were washed with PBST and incubated at RT for 1 h with PBST containing 1% BSA, and 5% FBS (blocking solution). Sections were then washed with PBST and incubated overnight at 4°C with the corresponding primary antibodies diluted in blocking solution. The next day, they were washed with PBST and incubated at RT for 2 h with the corresponding secondary antibodies diluted in blocking solution, washed again, incubated with DAPI (1:2000 in PBS) for 10 min at RT, and mounted in Superfrost^TM^Plus Adhesion Microscope Slides (Epredia, USA) with mowiol. In some cases, ThioS-flavin (ThioS) staining was performed right before starting the immunofluorescence protocol. Briefly, sections were washed with 50% ethanol (EtOH), incubated with 0.01% ThioS/EtOH for 10 min, and thoroughly washed with EtOH and then water. All sections were imaged and photographed using a Zeiss LSM900 upright confocal microscope (Zeiss, Germany) with a 10x/0.3 Plan-Apo air objective and the ZEN Blue 3.3 software (Zeiss, Germany). A minimum of 3 and a maximum of 8 slices were photographed for each sample. Image analysis was performed using custom macros with FIJI software (35).

#### 2.13. Hematoxylin-Eosin staining

Cryostat sections (15µm) mounted in Superfrost^TM^Plus Adhesion Microscope Slides (Epredia, USA) were allowed to thaw at RT and air-dried. Slides were rinsed in water for 5 min and dehydrated through a graded ethanol series (70%, 90%, and 100% EtOH; 5 min each), followed by rinses in water. Slides were stained with hematoxylin (Sigma-Aldrich) for 4 min, thoroughly washed with abundant water and later incubated with eosin (Sigma-Aldrich) for 3 min. Slides were thoroughly washed with water under the tap and then dehydrated with rinses in a sequential EtOH series and then left in 100% for 5 min. Slides were then cleared in xylene twice for 10 min and mounted with DPX mounting medium (Sigma-Aldrich). Images were analyzed and photographed using a Leica microscope (DM5000) equipped with digital camera (DFC350FX). Images were analyzed and processed with the ImageJ software.

#### 2.14. Statistical analysis

Statistical significance of differences between groups was calculated by 2-way ANOVA followed by Sedak’s test for multiple comparisons or by two-tailed, unpaired Student’s t-test or Mann-Whitney test. Differences were concluded significant for p values < 0.05. All statistical tests were performed using GraphPad Prism 8.0.1 (GraphPad Software, USA).

## 3. Results

### 3.1 Retro-orbital vs intraperitoneal mAb administration

In previous studies, we reported that α-SFRP1 (IgG1 subtype) can efficiently neutralize SFRP1-mediated ADAM10 inhibition. Furthermore, we demonstrated that the mAb reached the brain parenchyma of APP/PS1 mice 24 hours after administration through the retro-orbital (RO) sinus (17). This route of administration requires mouse anaesthetization, increasing animal stress (36) and represents a more challenging and time-consuming procedure than intraperitoneal (IP) inoculation. With the aim of improving animal welfare while facilitating the experimenter’s work, we first compared the kinetic and half-life of a biotinylated form of the α-SFRP1 (α-SFRP1^biot^; 100 µg) after either RO or IP administration into 4-8 months old homozygous APP/PS1 mice (APP/PS1^Hom^), when amyloid plaque deposits are already evident (17). α-SFRP1^biot^ levels were then measured in the serum and brain extracts of the injected mice, using a home-made enzyme-linked immunosorbent assay (ELISA). RO inoculation resulted in 3-fold increase in mAb serum levels relative to IP, at least during the first 6 hours post-injection (**Fig. S1A**) with a half-life of 9.17 hours. However, after 24 hours, both routes of administration showed comparable levels of α-SFRP1^biot^ in brain and serum (**Fig. S1A,B**).

These results suggest that RO administration enables a faster and more efficient distribution of the mAb, at least in the first few hours after inoculation, making it the route of choice for the following experiments.

### 3.2 α-SFRP1 is widely distributed across the mouse body, including the brain

We next analysed the mAb biodistribution across the mouse body and compared its behaviour with that of a biotinylated form of an unspecific and isotypic control antibody (IgG1^biot^), administered in 4 to 8-months old APP/PS1^Hom^ mice. Antibody levels were measured 24 hours later in various organs (serum, liver, kidney, brain). Comparable concentrations of both antibodies were detected in the bloodstream at 6- and 24-hours post-injection (**Fig. S2A**). However, α-SFRP1^biot^ seemed to accumulate in the kidney, where SFRP1 is expressed (37) as compared to IgG1^biot^, whereas equal levels of both antibodies reached the brain and liver (**Fig. S2B-D**).

To obtain a more accurate estimate of α-SFRP1 biodistribution, we inoculated Zirconium-89 (^89^Zr) radiolabeled antibodies (α-SFRP1^89Zr^ and IgG1^89Zr^; 100 µg) and then used positron emission tomography in combination with computed tomography (PET/CT) for longitudinal and whole-body tracing (2, 10, 24, 168 hours) of the antibodies in a group of 3-4 animals per treatment (**Fig. 1A**). IgG1^89Zr^ concentrated in the bladder at earlier time points, suggesting faster clearance from the vascular system as compared to α-SFRP1^89Zr^ (**Fig. 1G**). In fact, α-SFRP1^89Zr^ specifically accumulated in the brain, lungs, and heart up to 10 hours post-injection, decreasing to IgG1^89Zr^ comparable levels 24 hours after the inoculation (**Fig. 1B,E,F**). Surprisingly, we observed no significant differences in the concentrations of antibodies in the liver (**Fig. 1C**) or kidneys (**Fig. 1D**), in contrast to our previous results obtained with α-SFRP1^biot^ and IgG1^biot^.

**Figure 1.**
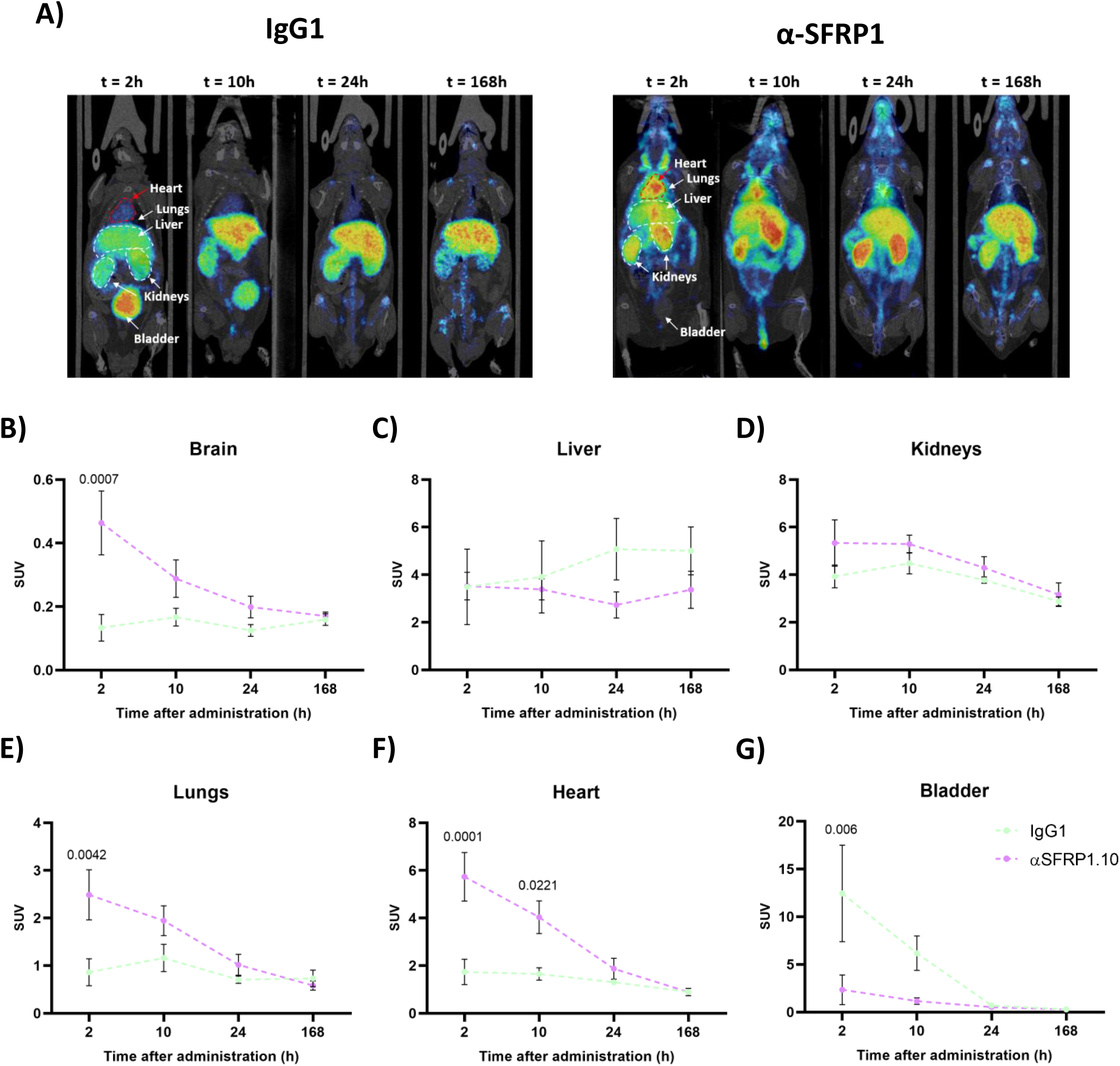
Whole-body PET–MRI reveals transient differences in systemic biodistribution of ^89^Zr-IgG1 and ^89^Zr-α-SFRP1.10 after intravenous administration. **(A)** Representative transverse whole-body PET-MRI maximum-intensity projection images acquired at 2, 10, 24, and 168 h after tail-vein administration of ^89^Zr-IgG1 (left) or ^89^Zr-α-SFRP1.10 (right). **(B-G)** *In vivo* quantification of ^89^Zr-IgG1 and ^89^Zr-α-SFRP1 overtime in **(B)** brain, **(C)** liver, **(D)** kidneys, **(E)** lungs, **(F)** heart, **(G)** and bladder. Data are presented as standardized uptake values (SUV) and shown as mean ± SEM. Statistical analysis were performed using two-way ANOVA followed by Sidak’s post-hoc correction for multiple comparisons.

α-SFRP1^89Zr^ accumulated throughout the brain when compared to IgG1^89Zr^ (**Fig. 2A**), with higher concentrations in the olfactory bulbs and entorhinal cortex, where SFRP1 is more abundantly distributed (37). In APP/PS1 mice, the cortex and the hippocampus are the brain areas with higher accumulation of amyloid and SFRP1 positive plaques (17,32). Both regions were specifically targeted by α-SFRP1^89Zr^, although antibody concentrations reached IgG1^89Zr^ comparable levels within the first 24 hours (**Fig. 2B,C**), mirroring whole brain observations.

**Figure 2.**
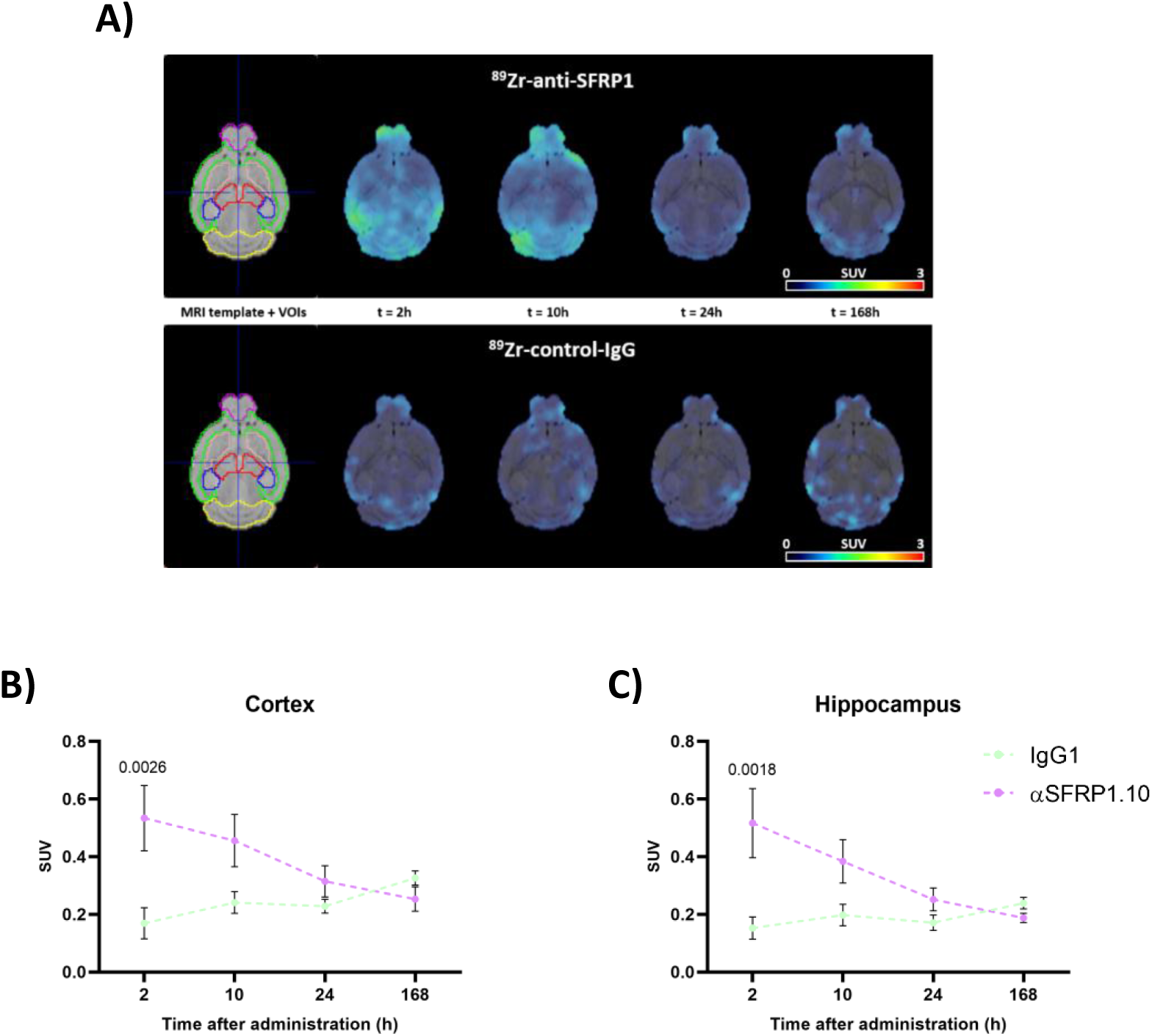
Regional brain PET–MRI analysis reveals transient differences in ^89^Zr-IgG1 and ^89^Zr-α-SFRP1 uptake after intravenous administration. **(A)** Representative transversal brain PET-MRI maximum-intensity projection images acquired at 2, 10, 24, and 168 h after intravenous tail administration of ^89^Zr-IgG1 (bottom) or ^89^Zr-α-SFRP1 (top). The left-most image corresponds to the mouse brain template used for the analysis. Volumes of interest include the olfactory bulb (purple), cerebral cortex (green), striatum (orange), hippocampus (blue), thalamus (red), and cerebellum (yellow). (B, C) *In vivo* quantification of ^89^Zr-IgG1 and ^89^Zr-α-SFRP1 over time in the brain regions of primary interest: **(B)** cortex and **(C)** hippocampus. Data are presented as standardized uptake values (SUV) and shown as mean ± SEM. Statistical analysis were performed using two-way ANOVA followed by Sidak’s post-hoc correction for multiple comparisons.

Together, these results show that the α-SFRP1 engages its target in different organs, reaching the brain, including the cortex and hippocampus of APP/PS1^Hom^ mice, where it could exert on-site beneficial effects mitigating AD progression. Notably, α-SFRP1 seems to be cleared from the brain within the first 24 hours after injection, allowing only a small time-window for its therapeutic potential.

### 3.3 α-SFRP1 administration has a reduced efficacy at later stages of the pathology

Amyloid-β plaques start to appear as early as 3 months in the cortex and hippocampus of APP/PS1^Hom^ mice (17) and they are followed by a marked astro- and microgliosis, neuronal loss, and cognitive impairment (32,38). We have previously treated APP/PS1^Hom^ mice with α-SFRP1 at 2 months of age, before the onset of the disease, observing a marked reduction of amyloid burden and related pathological hallmarks (17). However, diagnosis of dementia-associated diseases occurs when the brain of the patients has undergone mayor morphological alterations (39), hindering the effects of the attempted treatments (40,41).

Thus, we tested whether α-SFRP1 could prove beneficial at later stages of the disease. To answer this question, we performed weekly injections of α-SFRP1 or IgG1 (100 µg, RO) in 9-month old APP/PS1^Hom^ mice (**Fig. S3A**), once the AD-like phenotype is fully developed, including cognitive deficits (42). After 3 months of treatment, animals were sacrificed (12 months of age) and their brain analysed. First, we employed a previously established SFRP1-specific and highly sensitive ELISA (17) to determine the levels of SFRP1 present in one brain hemisphere RIPA-soluble fractions, finding no significant changes in SFRP1 concentration between control and α-SFRP1 treated animals (**Fig. S3B**). We next investigated the effects of the treatment on amyloid burden, neuronal damage, and gliotic response by immunostaining cortical and hippocampal slices with α-Aβ42 (H31L21), α-GFAP, and α-IBA1. We observed 33% decrease in the number of Aβ^+^ plaques, although this decrease did not reach statistical significance, and no difference in Aβ^+^ plaque size, in the cortex of α-SFRP1-treated animals. Notably, α-SFRP1 had no apparent effects on the Aβ plaque deposition in the hippocampus (**Fig. S3D**). Furthermore, the levels of astrocytic and microglial activation, measured as GFAP and IBA1 fluorescence intensity, remained unaltered between the two experimental groups (**Fig. S3E,F**). We also observed no changes in the percentage of different blood cell populations, suggesting no apparent adverse peripheral effects of the treatment (**Fig S4**).

Similar results were obtained when we treated 4-month old APP/PS1^Hom^ mice with weekly mAb administration (100 µg, RO) for 5 months (**Fig. 3A**). No significant changes were observed in SFRP1 levels present in the RIPA-soluble brain homogenates (**Fig. 3B**) or in the amyloid burden (**Fig. 3D**), dystrophic neuron load (**Fig. 3E**), or astro- and microgliosis (**Fig. 3F,G**) of the cortex and hippocampus of these mice. These mice showed no detectable defects in the overall structure of their liver and kidneys (**Fig. S5).**

**Figure 3.**
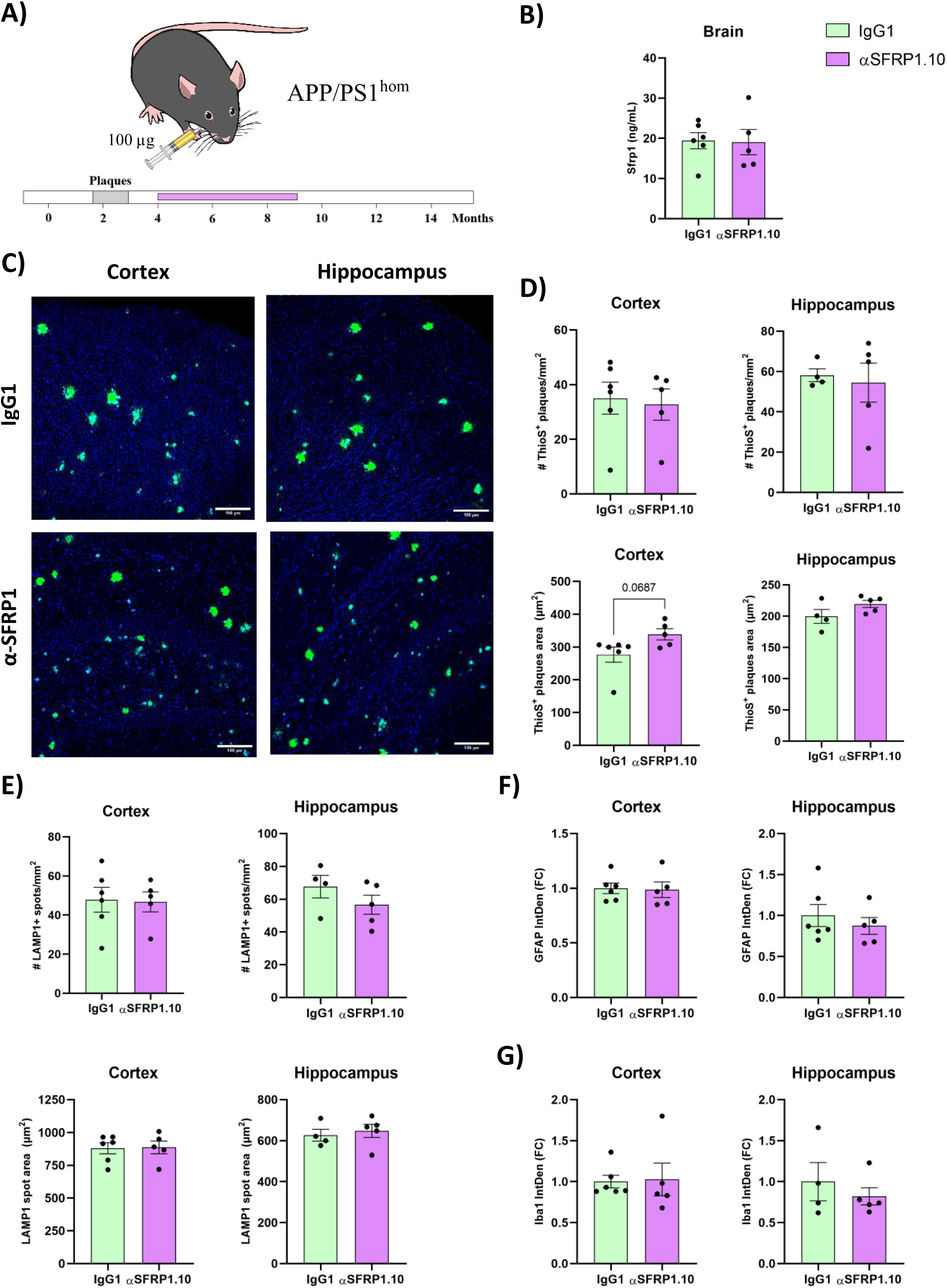
Long-term α-SFRP1 treatment does not significantly modify AD pathophysiology in adult APP/PS1 mice. (**A**) Schematic of the experimental design. 4-month-old homozygous APP/PS1 mice received weekly retro-orbital injections of IgG1 or α-SFRP1.10 (100 µg) for 5 months; analyses in panels (**B–G**) were performed at 9 months of age. The time diagram indicates the expected age at which these mice initiate amyloid-β plaque accumulation. Illustration from NIAID NIH BioArt Source (https://bioart.niaid.nih.gov/). (**B**) ELISA quantification of SFRP1 levels in brain RIPA-soluble homogenates. (**C**) Representative confocal images of cortices (top) and hippocampi (bottom) from IgG1-treated (left) and α-SFRP1.10-treated (right) mice stained with Thioflavin S (ThioS) to label amyloid plaque cores. Scale bar: 100 µm. Quantification of the number (top) and mean plaque area (bottom) of (**D**) ThioS+ plaques and (**E**) LAMP1-positive puncta in the cortex (left) and hippocampus (right). Quantification of **(F)** GFAP and **(G)** Iba1 fluorescence intensity in the cortex (left) and hippocampus (right), expressed as fold change relative to the mean value of the IgG1 group. Quantification analyses were performed for individual mice, averaging the results of 3-8 slices per mouse. Data are presented as mean ± SEM. Statistical analyses were performed using Student’s t test.

These results indicate that α-SFRP1, at the applied concentration, has little effect when the pathological alterations are well established in the brain of APP/PS1^Hom^ mice. We thus tested whether a higher dose of α-SFRP1 would be necessary to show beneficial effects in APP/PS1^Hom^ mice in these conditions. To this end, we used 4-month old APP/PS1^Hom^ mice and inoculated them weekly with α-SFRP1 or IgG1 (200 µg, RO) (**Fig. 4A**). The experiment was terminated after 2 months due to the low survival rate (43%) in the α-SFRP1-treated group (**Fig. 4B**). Surviving α-SFRP1-treated animals showed a 50% decrease in SFRP1 brain concentration as compared with IgG1 injected mice (**Fig. 4C**) and their number of ThioS^+^ plaques was reduced by 46 and 57% in the cortex and hippocampus, respectively (**Fig. 4E**). The amelioration of Aβ load was paralleled by a reduction in the number of LAMP1^+^ spots surrounding the plaques in the cortex (37%) and hippocampus (39%; **Fig. 4F**). In both cases, the area covered by amyloid plaques and dystrophic neurons remained unchanged (**Fig. 4E,F**). Although not statistically significant, additional differences were also noted, particularly in microglial activation, with a decreased IBA1^+^ signal intensity in the cortex (35%) and hippocampus (49%) of the surviving α-SFRP1 treated mice (**Fig. 4H**). Conversely, the astrogliotic response was unaffected in the cortex, while it showed a non-significant 40% increase of GFAP intensity in the hippocampus (**Fig. 4G**).

**Figure 4.**
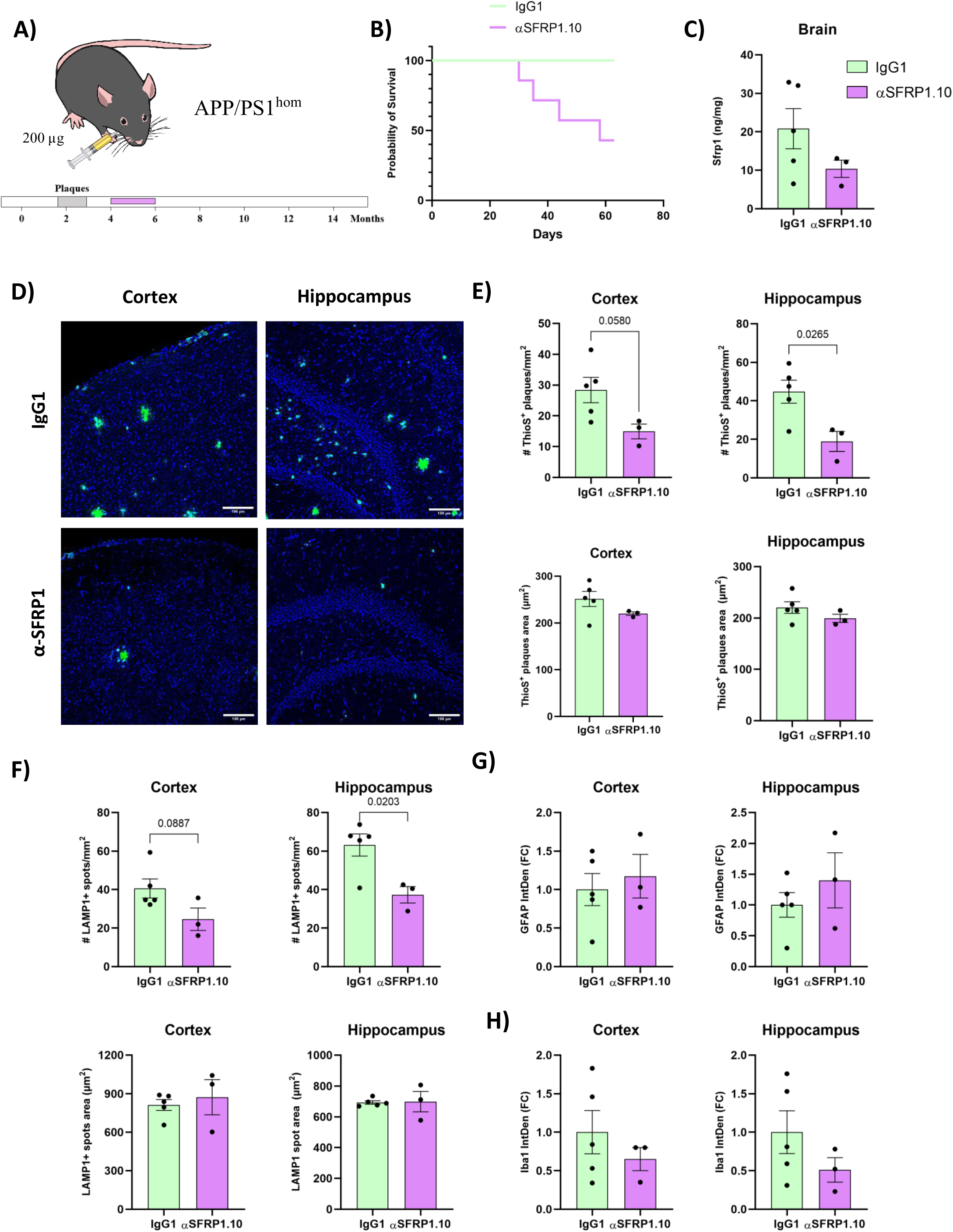
High-dose α-SFRP1 treatment reduces amyloid burden in adult APP/PS1 mice. (**A**) Schematic of the experimental design. 4-month-old homozygous APP/PS1 mice received weekly retro-orbital injections of IgG1 or α-SFRP1.10 (200 µg) for 2 months; analyses in panels (**B–H**) were performed at 6 months of age. The time diagram indicates the expected age at which these mice initiate amyloid-β plaque accumulation. Illustration from NIAID NIH BioArt Source (https://bioart.niaid.nih.gov/). (**B**) Kaplan–Meier curves showing mouse survival during the treatment period; each vertical step represents a death event. (**C**) ELISA quantification of SFRP1 levels in brain RIPA-soluble homogenates. (**D**) Representative confocal images of cortices (top) and hippocampi (bottom) from IgG1-treated (left) and α-SFRP1.10-treated (right) mice stained with Thioflavin S (ThioS) to label amyloid plaque cores. Scale bar: 100 µm. Quantification of the number (top) and mean plaque area (bottom) of (**E**) ThioS+ plaques and (**F**) LAMP1+ puncta in the cortex (left) and hippocampus (right). Quantification of **(G)** GFAP and **(H)** Iba1 fluorescence intensity in the cortex (left) and hippocampus (right), expressed as fold change relative to the mean value of the IgG1 group. Quantification analyses were performed for individual mice, averaging the results of 3-8 slices per mouse. Data are presented as mean ± SEM. Statistical analyses were performed using Student’s t test.

Taken together these results indicate that higher doses of α-SFRP1 are required to alleviate AD traits in mice when brain morphological alterations are well established. Unfortunately, this dose is associated with an increased mortality, the causes of which remained unexplored.

### 3.4 Chemical inhibition of SFRP1 does not alleviate AD at intermediate stages of the disease

There is substantial evidence that brain-targeted immunotherapies highly increase the risk of patients from suffering a type of brain edemas and hemorrhages known as Amyloid-Related Imaging Abnormalities (ARIA) (43–47), which may represent the reason behind the premature death in mice treated with higher α-SFRP1 doses. With a view to circumvent this problem, while still targeting SFRP1 in the brain, we tested WAY-316606 (WAY), a specific SFRP1 inhibitor, for its potential efficacy. WAY has been previously shown to decrease Aβ production in AD patient-derived organoids (48) and to improve experimentally induced osteoporosis *in vivo* (49), with no apparent toxic side effects.

To test this compound, we employed heterozygous APP/PS1 mice (APP/PS1^Het^), which show a slower progression of the AD-like phenotype, starting to show Aβ plaque accumulation at 6 months of age (32). Therefore, we treated 8-month old APP/PS1^Het^ mice with RO weekly injections of either 1 µM WAY or 2% DMSO, as a control, for 2 months (**Fig. 5A**). We observed a non-significant 34% reduction in the levels of SFRP1 in WAY-injected mice brains (**Fig. 5B**). However, no apparent differences were found in the number or size of ThioS^+^ amyloid plaques or LAMP1^+^ dystrophic neuron spots (**Fig. 5D,E**).

**Figure 5.**
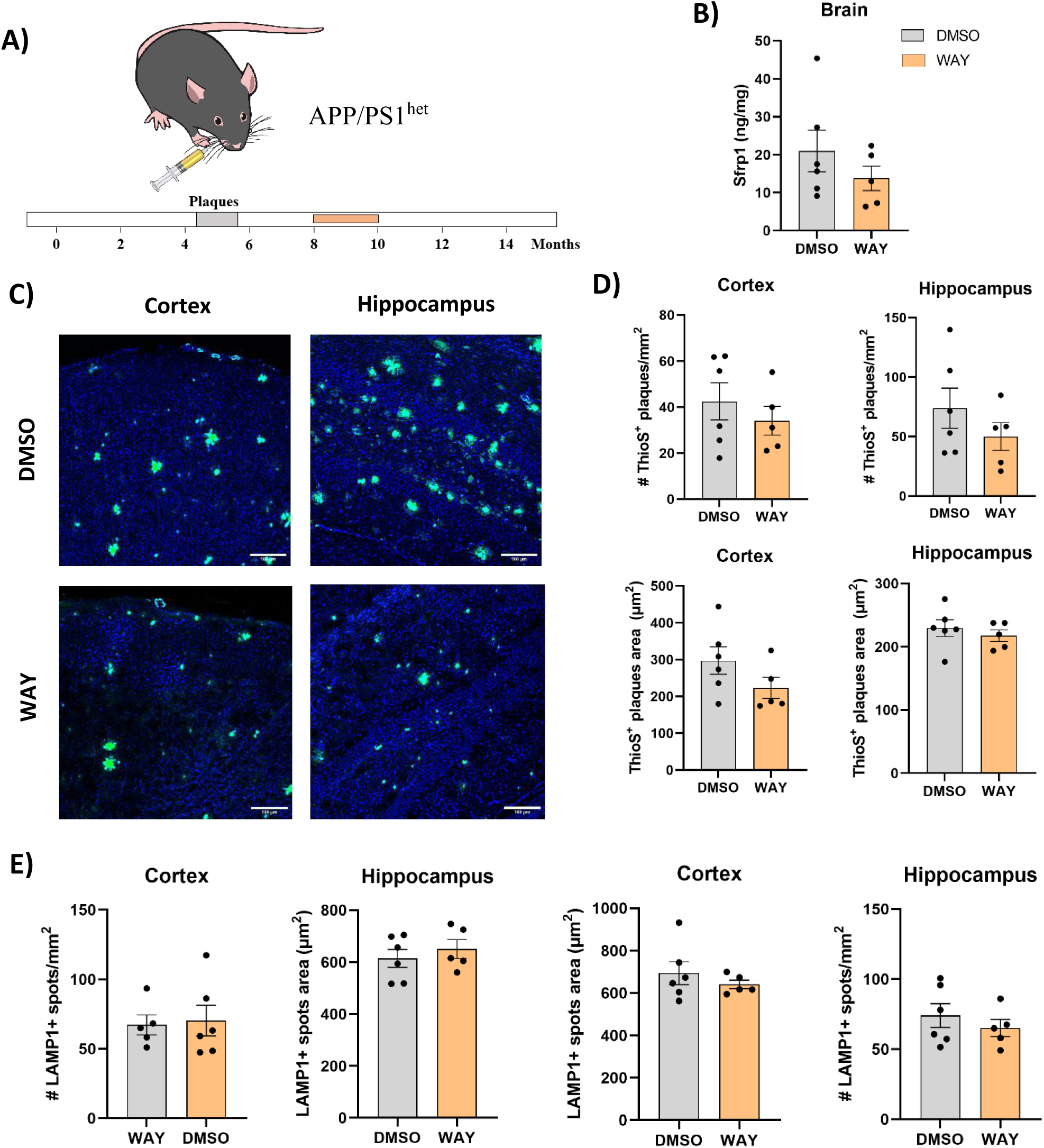
Pharmacological inhibition of SFRP1 do not significantly modify AD related pathology in adult APP/PS1 mice. (**A**) Schematic of the experimental design. 8-month-old heterozygous APP/PS1 mice received weekly retro-orbital injections of WAY-316606 (1 µM) or vehicle (2% DMSO) for 2 months; analyses in panels (**B–D**) were performed at 10 months of age. The time diagram indicates the expected age at which these mice initiate amyloid-β plaque accumulation. Illustration from NIAID NIH BioArt Source (https://bioart.niaid.nih.gov/). (**B**) ELISA quantification of SFRP1 levels in brain RIPA-soluble homogenates. (**C**) Representative confocal images of cortices (top) and hippocampi (bottom) from vehicle-treated (left), WAY-316606-treated (center) mice stained with ThioS to label amyloid plaque cores. Scale bar: 100 µm. Quantification of number (top) and mean plaque area (bottom) of (**D**) ThioS+ plaque and LAMP1+ puncta (**E**) in the cortex (left) and hippocampus (right). Quantification analyses were performed for individual mice, averaging the results of 3-8 slices per mouse. Data are presented as mean ± SEM. Statistical analyses were performed using Student’s t test.

Taken together, these data indicate that the α-SFRP1 mAb may represent a better tool to target SFRP1 in AD conditions, although optimization of its dose and delivery to the brain is needed.

## 4. Discussion

AD is characterized by a prolonged preclinical phase during which subtle molecular and cellular alterations progressively damage synaptic integrity, disrupt neuronal plasticity, and promote amyloid-driven neurodegeneration (5,50–52). By the time cognitive symptoms emerge, the large number of well-known morphological alterations are already fully settled, greatly limiting the efficacy of disease modifying interventions. This temporal disconnection between disease onset and clinical diagnosis remains one of the major challenges for successful AD treatment (39,41,53,54). This together with the complexity of AD onset, underscore the need of selecting targets with pleiotropic functions, early appearance and efficacy beyond the prodromic phase. A recent proteomic study in cohorts of AD patients has confirmed the accumulation of SFRP1 at prodromal diseases stages (21), whereas functional studies have demonstrated its involvement in several key processes that precede overt neurodegeneration (29,30,48,55). These observations strongly suggest that its neutralization may have multiple beneficial effects to slow down AD progression. The present study extends these conclusions and supports the potential therapeutic value of mAb-mediated SFRP1 neutralization, even after full establishment of AD pathology. However, it also shows that its efficacy is strongly constrained by limited brain targeting and higher dose requirement that associates with an increased animal mortality.

We have previously shown that early SFRP1 neutralization using low doses of α-SFRP1 reduces plaque burden, neuroinflammation and preserves synaptic plasticity in APP/PS1 mice (17), leaving the pharmacokinetic, biodistribution properties, and therapeutic window of the mAb largely unexplored. In this study, we show that RO administration leads to significantly higher circulating α-SFRP1 levels as compared with IP delivery. Even though IP injections are more amenable and less stressing for preclinical studies in mice, IV delivery is commonly used in human antibody-based therapies, thus offering greater translational relevance (56).

Using biotinylated and ^89^Zr-radiolabeled antibodies combined with PET/CT imaging, we further demonstrate that IV administered α-SFRP1 engages its target systemically, accumulates in different organs, and reaches the brain, including AD affected regions such as the cortex and hippocampus. Nonetheless, brain exposure was markedly low relative to peripheral organs such as liver and kidney, and declined rapidly within 24 hours. This observation aligns with extensive literature showing that under physiological conditions only a small fraction (∼0.1%) of circulating IgG crosses the intact blood–brain barrier (BBB) (57). BBB integrity is partially compromised in AD, particularly in regions affected by amyloid pathology (58,59). Consistent with this observation, the detectable hippocampal accumulation observed in our study is biologically plausible and comparable to what has been reported for other therapeutic antibodies targeting amyloid-β in transgenic mouse models (46,60). Nevertheless, we found ten-fold less concentration in the brain than in the kidney or the liver, indicating that mAb penetration remains limited even in AD conditions and still represents a major obstacle for CNS immunotherapies. It is important to note that IV inoculation of drugs was consistently performed through the RO sinus, with the exception of the ^89^Zr-radiolabeled antibodies, which were administered through de tail vein. Given that both of these routes have been shown to provide comparable absorption and pharmacokinetic activity for therapeutic antibodies (61), they were considered equivalent for the purpose of this study.

Besides this restricted brain availability, the therapeutic efficacy of α-SFRP1 proved highly dependent on disease stage. When treatment was initiated in mice with established plaque pathology (4–9 months of age), standard antibody dosing failed to significantly reduce SFRP1 concentration in the brain. This may reflect that SFRP1 accumulates in the plaques and cannot be neutralized, further becoming resistant to degradation, as shown in a recent study (27). This persistent SFRP1 presence may explain also why amyloid burden, neuritic dystrophy, or gliosis are unaffected by the mAb administration.

Only upon dose escalation did surviving animals show reductions in plaque number and dystrophic neurites, suggesting that SFRP1 remains mechanistically involved even at later stages, but cannot be effectively neutralized at conventional doses due to limited CNS exposure. This underscores a key challenge shared across neurodegenerative immunotherapies: sufficient target engagement in the brain often requires antibody concentrations that are difficult to achieve safely through systemic administration. In fact, similar dose-limiting effects have been described for other antibody-based therapies in ND models. For example, higher doses required to achieve clinical efficacy with antibodies such as Aducanumab and Lecanemab have been associated with significant adverse effects (46,62).

The increased mortality observed at higher antibody doses further highlights the narrow therapeutic margin for CNS-directed immunotherapy in AD. Although the causes of death were not directly investigated in this study, one plausible explanation relates to vascular complications reminiscent of Amyloid-Related Imaging Abnormalities (ARIA), which arise from antibody-mediated clearance of vascular amyloid deposits and are frequently associated with Aβ-targeting antibodies such as Aducanumab, Lecanemab, and Donanemab (43,45). While α-SFRP1 does not directly target Aβ, its ability to modulate plaque-associated pathology may similarly influence vascular amyloid dynamics or inflammatory responses, particularly when administered at supratherapeutic doses.

The commercial availability of small-molecule SFRP1 inhibitors, such as WAY-316606, prompted us its trail as an alternative to α-SFRP1. However, this pharmacological modulation was insufficient to reverse pathology once amyloid deposition is established. Although WAY has been reported to reduce Aβ production in patient-derived organoids (48), its administration in intermediate-stage APP/PS1^Het^ mice produced only modest and non-significant reductions in brain SFRP1 levels, without measurable effects on plaque burden or neuritic dystrophy.

Collectively, our study defines critical pharmacodynamic and temporal constraints for targeting SFRP1 in AD. While SFRP1 neutralization remains a valid strategy, its therapeutic benefit is greatest at early disease stages, before widespread plaque deposition, glial activation, and synaptic degeneration. These findings parallel the broader field of AD therapeutics, where clinical benefit from anti-Aβ antibodies is similarly restricted to prodromal or very early symptomatic populations. Thus, SFRP1-directed therapies may ultimately prove most effective as part of an early intervention strategy, potentially combined with biomarkers for patient stratification and improved brain-targeted delivery platforms.

Current AD therapies either aim to alleviate symptoms, including, for example, acetylcholinesterase inhibitors and NMDA receptor antagonists, or are disease-modifying compounds, such as the recently approved anti-amyloid mAb drugs Aducanumab, Lecanemab, and Donanemab (16,63). Systematic reviews of the current literature and regulatory reports indicate that at least Lecanemab and Donanemab, when administered to patients at early stage of the disease can slow disease progression by approximately 30%, but are associated with side effects such as ARIAs and a rather heavy following-up protocol (43,53). Our preclinical analysis in mice provides a comprehensive evaluation of α-SFRP1 pharmacodynamics and stage-dependent efficacy *in vivo*, suggesting that this approach could be used in alternative or in combination with current treatments. However, additional efforts are needed to optimize its dose and brain delivery. Strategies such as BBB shuttle engineering, Fc receptor-mediated transcytosis, or encapsulation into brain-targeted lipid vesicles are possible options to substantially expand the therapeutic window of α-SFRP1. Combinatorial strategies, including SFRP1 neutralization with the currently approved Lecanemab or Donanemab may provide synergistic benefit by simultaneously facilitating plaque clearance (64) and restoring non-amyloidogenic APP processing (17), while reducing neuroinflammation (29) and preventing synaptic alterations (30).

## Data statement

All data generated or analysed during this study are included in this published article and its supplementary materials.

## Declaration of the use of generative AI and AI-assisted technologies in scientific writing and in figures, images and artwork

Generative AI was not used in the design, analysis or writing of this work

## Ethical statement

All experiments were performed according to procedures approved by the ethical committees of the CBM, the Bioethics sub-committee of Consejo Superior de Investigaciones Científica (CSIC, Madrid, Spain) and the Comunidad Autónoma de Madrid under the protocol approval number PROEX 092.6/21; RD 53/2013 or the Ethical Committee of Centro de Investigación Cooperativa en Biomateriales (CIC biomaGUNE) and authorized by the local Diputación Foral de Gipuzkua under the protocol approval number PRO-AE-SS-212 in case of radiolabeled antibodies administration.

## Authors’ contributions

PM and MMB performed all experiments with the help of MJMB. PM and MMB analyzed and interpreted the data with PB. IM and MD generated, purified and verified the activity of the mAb. PM drafted the manuscript and PB produced the final version. All authors read and approved the final manuscript. PB supervised the work and obtained funds.

## Acknowledgements

The authors thank CIC BiomaGUNE for performing the PET/CT biodistribution studies of 89Zr-labeled antibodies and for their technical assistance. We are in debt with the Advanced Light Microscopy and Preclinical Biomedicine facilities of the CBM for their expert and invaluable support.

---

**Figure S1.**
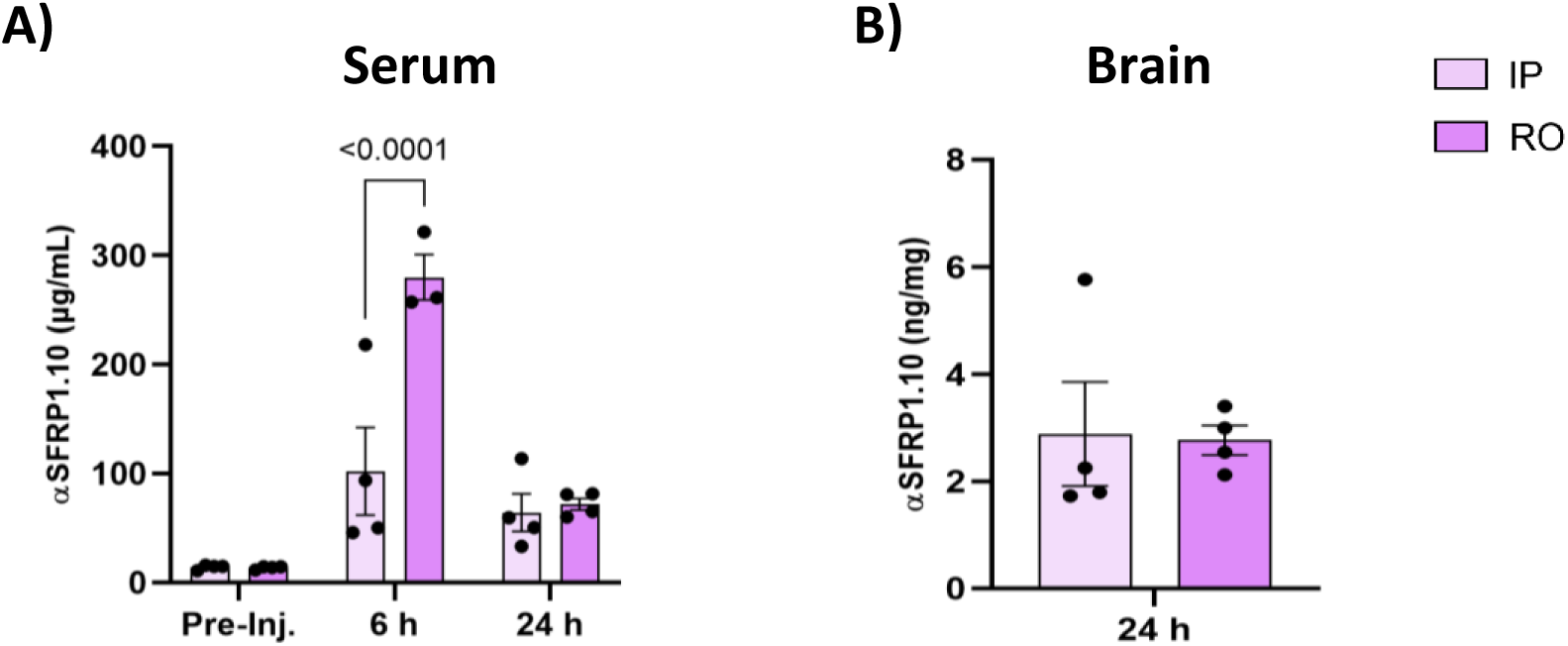
Retro-orbital injection results in higher short-term vascular exposure to α-SFRP1 than intraperitoneal administration. ELISA quantification of biotinylated α-SFRP1 mAb levels in **(A)** serum and **(B)** brain RIPA-soluble homogenates from 4-8 month-old APP/PS1 mice. Samples were collected 6 and 24 h after a single 100 µg intraperitoneal or retro-orbital injection of α-SFRP1. “Pre-Inj.” Indicates mAb levels before the injection (expected to be zero). Data are presented as mean ± SEM. Statistical analyses were performed using **(A)** two-way ANOVA followed by Sidak’s post hoc test for multiple comparisons or **(B)** Mann-Whitney test.

**Figure S2.**
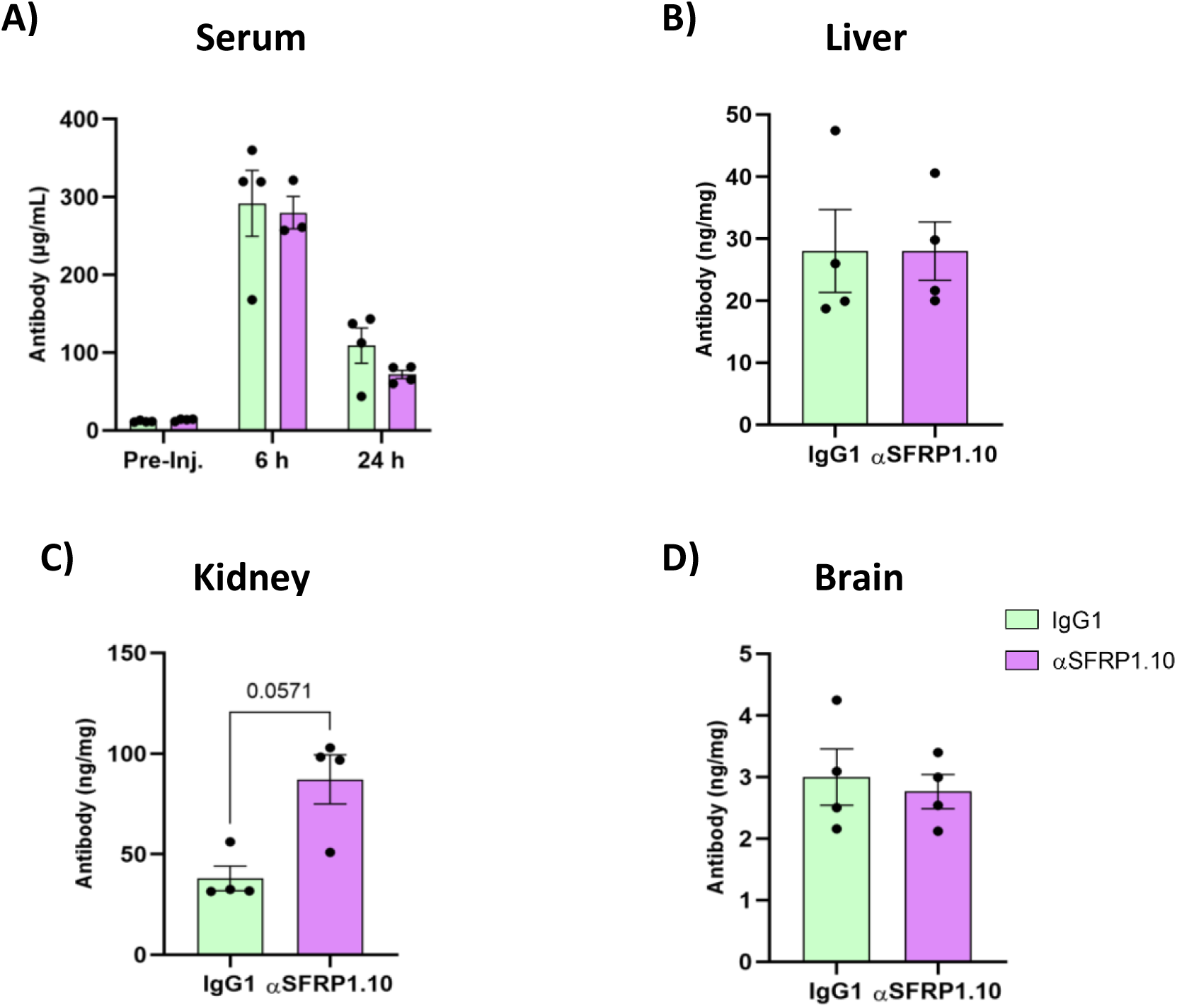
Biotinylated IgG1 and α-SFRP1 show a similar organ distribution 24 h after retro-orbital injection. ELISA quantification of biotinylated IgG1 and α-SFRP1 mAb levels in **(A)** serum, **(B)** liver, **(C)** kidney, and **(D)** brain RIPA-soluble homogenates from 4-8 month-old APP/PS1 mice 24 h after a single 100 µg RO injection of either mAb. Data are presented as mean ± SEM. Statistical analysis were performed using **(A)** two-way ANOVA followed by Sidak’s post-hoc correction for multiple comparisons, **(B, D)** Student’s *t*-test or **(C)** Mann-Whitney test.

**Figure S3.**
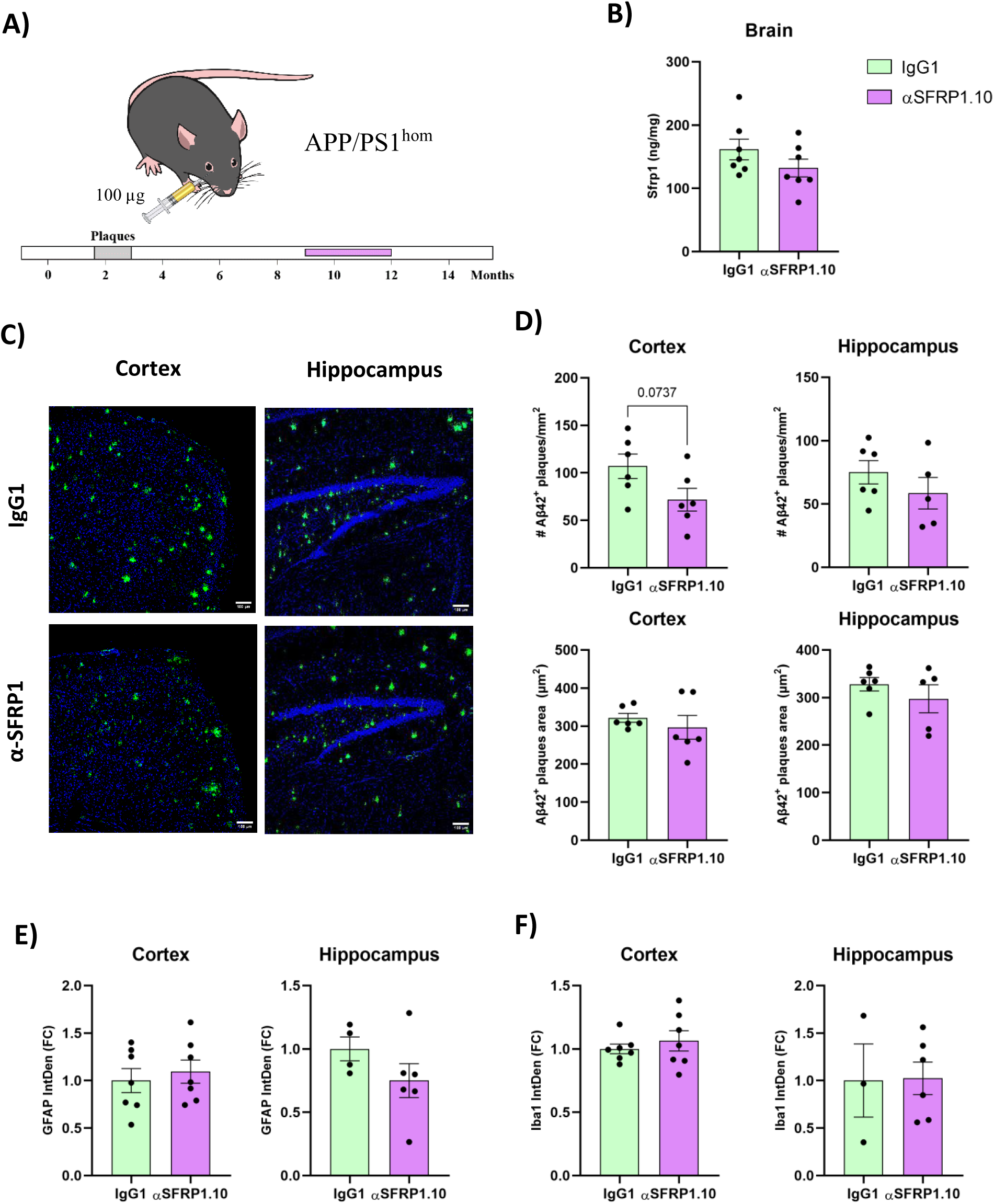
α-SFRP1 treatment does not significantly modify AD–related pathology in aged APP/PS1 mice. **(A)** Schematic representation of the experimental design: 9-month-old APP/PS1 homozygous mice received a weekly RO injection of IgG1 or α-SFRP1 (100 µg) for 3 months; analysis in panels (**B-F)** were performed at 12-months of age. The time diagram indicates the expected age at which these mice initiate amyloid-β plaque accumulation. Illustration from NIAID NIH BioArt Source (https://bioart.niaid.nih.gov/). **(B)** ELISA quantification of of SFRP1 levels in brain RIPA-soluble homogenates. **(C)** Representative confocal images of cortices (top) and hippocampi (bottom) from IgG1 (left) and α-SFRP1 (right) treated mice stained for Aβ42 as a marker of amyloid plaques. Scale bar: 100 µm. **(D)** Quantification of Aβ42^+^ plaques number (top) and mean plaque area (bottom) in the cortex (left) and hippocampus (right). **(E)** Quantification of GFAP and **(F)** Iba1 fluorescence intensity in the cortex (left) and hippocampus (right), expressed as fold change relative to the mean value of the IgG1 group. Quantification analyses were performed for individual mice, averaging the results of 3-8 slices per mouse. Data are presented as mean ± SEM. Statistical analysis were performed using Student’s t-test.

**Figure S4.**
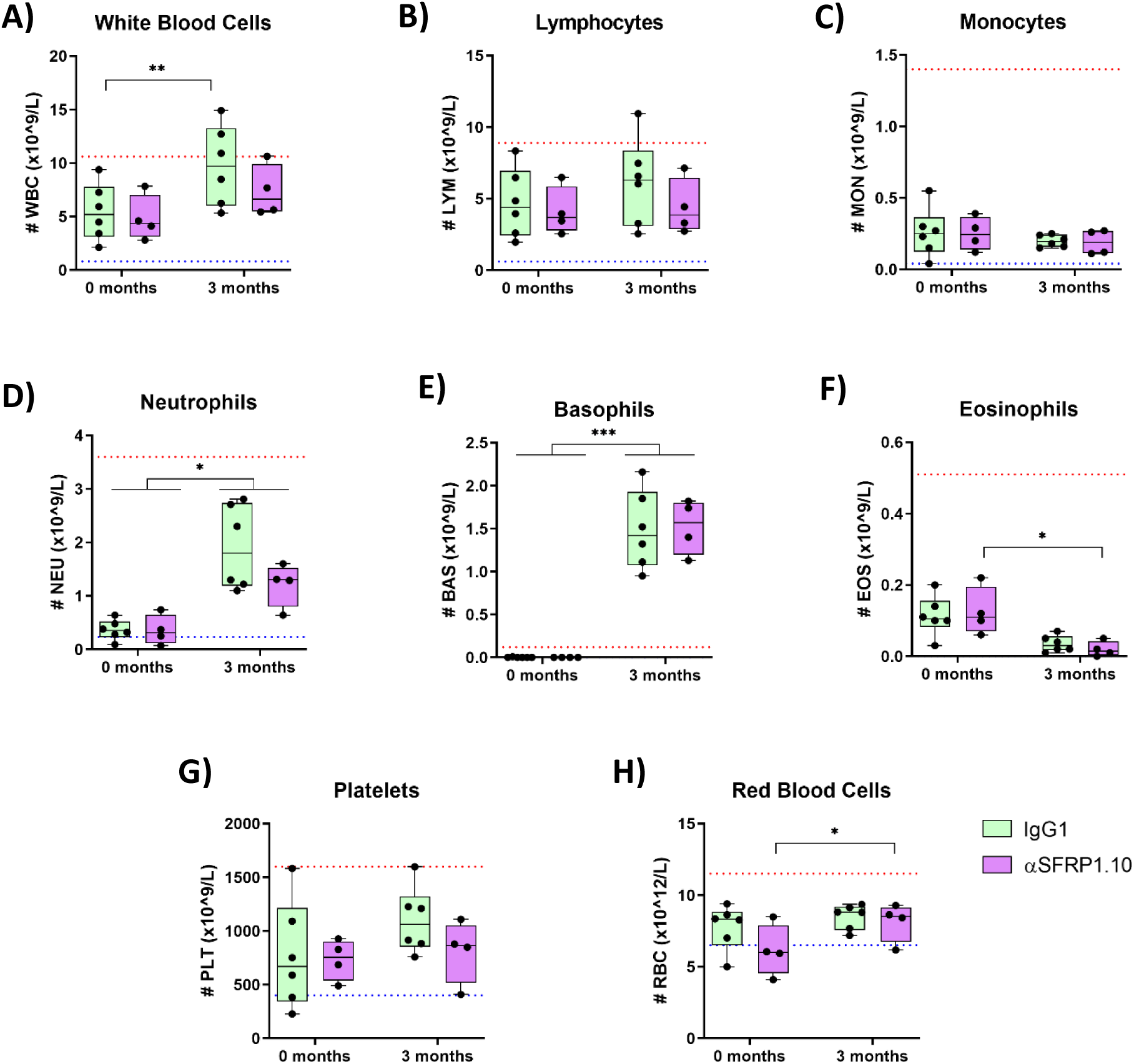
α-SFRP1 treatment does not affect blood cell populations count. Hematology analyzer (HT5) counts of whole blood cell populations in 12 month-old APP/PS1 homozygous mice. Mice were treated weekly with RO injections of IgG1 or α-SFRP1 (100 µg) for 3 months. Different blood cell types were analyzed at the beginning (0 months) and the end (3 months) of the treatment: **(A)** White blood cells, **(B)** lymphocytes, **(C)** monocytes, **(D)** neutrophils, **(E)** basophils, **(F)** eosinophils, **(G)** platelets, and **(H)** red blood cells. Dotted red (top) and blue (bottom) lines in each graph represent the reference range of values for healthy wild-type mice. Statistical analyses were performed using two-way ANOVA followed by Sidak’s post-hoc correction for multiple comparisons.

**Figure S5.**
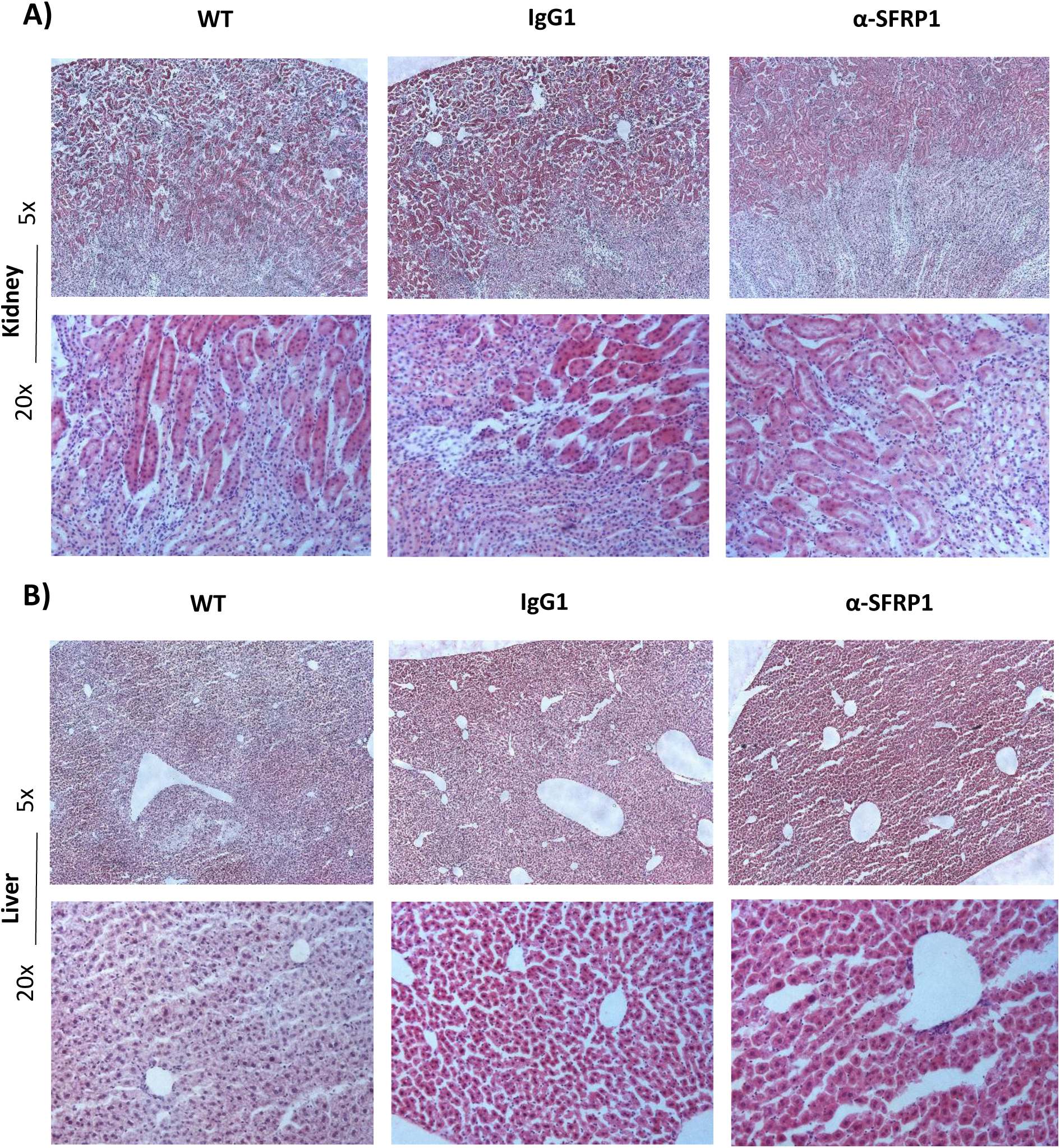
α-SFRP1 treatment does not affect overall liver and kidney structure. Representative images of hematoxylin/eosin stainings of the **(A)** kidney and **(B)** liver from 9 month-old mice. Images are shown at 5x (top) and 20x (bottom) magnifications for wild-type mice (left) and APP/PS1 homozygous mice treated with either IgG1 (middle) or α-SFRP1 (right). Mice received weekly retro-orbital injections of either antibody (100 µg) for 5 months. A minimum of 4 mice per group and 6-7 stained slices per mice were visually inspected and showed no apparent abnormalities.

